# Binding From both sides: TolR and full-length OmpA bind and maintain the local structure of the *E. coli* cell wall

**DOI:** 10.1101/409466

**Authors:** Alister T. Boags, Firdaus Samsudin, Syma Khalid

**Affiliations:** School of Chemistry, University of Southampton, Highfield, Southampton SO17 1BJ, UK

## Abstract

We present a molecular modeling and simulation study of the of the *E. coli* cell envelope, with a particular focus on the role of TolR, a native protein of the *E. coli* inner membrane in interactions with the cell wall. TolR has been proposed to bind to peptidoglycan, but the only structure of this protein thus far is in a conformation in which the putative peptidoglycan binding domain is not accessible. We show that a model of the extended conformation of the protein in which this domain is exposed, binds peptidoglycan largely through electrostatic interactions. We show that non-covalent interactions of TolR and OmpA with the cell wall, from the inner membrane and outer membrane sides respectively, maintain the position of the cell wall even in the absence of Braun’s lipoprotein. When OmpA is truncated to remove the peptidoglycan binding domain, TolR is able to pull the cell wall down towards the inner membrane. The charged residues that mediate the cell-wall interactions of TolR in our simulations, are conserved across a number of species of Gram-negative bacteria.

## INTRODUCTION

Gram-negative bacteria such as *E. coli* have a complex cell envelope, which protects the cell and controls influx/efflux of molecular species to ensure the normal functioning of the cell (Nikaido 2003). The cell envelope contains an aqueous region known as the periplasm, which is sandwiched between an asymmetric outer-membrane (OM) and a symmetric inner membrane (IM). Contained within the periplasm is the cell wall, which is composed of a sugar-peptide polymer known as peptidoglycan (PGN) (Vollmer and Bertsche 2008). The periplasm is host to many different proteins that are essential for the healthy growth and proliferation of Gramnegative bacteria. These proteins are known to be freely moving, associated with the inner or outer membranes, or bound to the cell wall. The interactions of these proteins with each other both (i) laterally, in other words within one membrane or the periplasm and (ii) across regions, e.g. extending from one membrane to the periplasm are important in maintaining the structural integrity and correct functioning of the cell envelope.

A number of different proteins have been shown to play a role in cross region interactions, and others have been hypothesized to do so. Braun’s lipoprotein (BLP, also known as “Lpp” and “murein lipoprotein”) is an abundant protein that is lipidated at its N-terminal domain, which anchors it to the outer membrane (Braun 1975). It is the only known protein in *E. coli* to be covalently attached to the PGN of the cell wall. It exists in two states; ~33% of the lipoprotein is covalently bound to the cell wall *via* a peptide bond, and ~66% is free in the periplasm. BLP is proposed to have a primarily structural function, essentially acting as a staple between the OM and PGN which serves to maintain the required distance between the cell wall and the OM (Miller and Salama 2018). Cells that lack BLP or that have reduced amounts of BLP are viable, but they have been shown to release OM vesicles at a higher rate than normal and also suffer from cellular leakage (Schwechheimer, Kulp et al. 2014, Asmar and Collet 2018). Non-covalent interactions between the cell wall and the outer membrane are mediated through proteins such as PAL and OmpA (Parsons, Lin et al. 2006, Park, Lee et al. 2012). The latter is composed of two domains, the N terminal domain which is an eight stranded beta barrel, this is connected via a flexible linker to the soluble C-terminal domain which contains the peptidoglycan binding region (Carpenter, Khalid et al. 2007, Marcoux, Politis et al. 2014). We have previously shown that OmpA in its dimeric form can extend its linker region such that the C-terminal is able to form long-lasting interactions with peptidoglycan even in the absence of BLP, while BLP facilitates peptidoglycan binding of the monomer (Samsudin, Boags et al. 2017). We showed that BLP can tilt within the periplasm to provide some variation in the PGN-outer membrane distance.

Interactions of inner membrane proteins with the cell wall are less well understood at the molecular level than their outer membrane counterparts. TolR is an inner membrane protein from the Tol family that is proposed to interact with the cell wall in a similar manner to OmpA. While binding of TolR to peptidoglycan has been demonstrated, the X-ray structure of TolR from *E. coli* is of the protein in its compact form, in which the putative PGN-binding domain is not surface exposed (Wojdyla, Cutts et al. 2015). Based on the X-ray structure of the closed state, biophysical and computational studies of TolR, Kleanthous and co-workers proposed a large-scale PMF-dependent conformational rearrangement in which extension of the TolR linker enables the protein to contact the cell wall and exposure of the PGN-binding domain enables it to bind PGN in a similar manner to the structural alterations proposed for the bacterial flagellar protein, MotB (Wojdyla, Cutts et al. 2015). The model of the protein in this conformation was termed the ‘open state’. The hypothesis of large-scale rearrangement is difficult to test experimentally in the absence of structures of the different states of the proteins. However simulations offer a route to predict the behaviour of the model under different scenarios.

To test the model of the open state of TolR and compare the PGN binding mode with that of OmpA, in the following we present an atomistic molecular dynamics and modelling study of TolR, OmpA, BLP in a model of the cell envelope that includes both membranes and the cell wall. We show that the model of the open state of TolR binds PGN primarily through electrostatic interactions, whereas the closed state does not bind PGN. In the presence of full-length OmpA dimers in the outer membrane and open state TolR in the inner membrane, the location of the cell wall is maintained between these proteins. The binding of both proteins to the cell wall also alleviates local surface distortion that are observed when only one out of OmpA or TolR is bound. In contrast if OmpA is truncated to its N-terminal domain and BLP is added to the system, then the TolR linker is able to contract and in doing so, ‘pulls’ the cell wall down towards the inner membrane until the BLP is fully stretched and further movement is not possible. This is strong indication that the strength of interaction with the cell wall of TolR and full length OmpA is similar.

## RESULTS

For ease of interpretation of the results the simulations described below are summarized in Table 1. The simulations of TolR and OmpA are performed with a monolayered cell wall. The reason for this is from test simulations of 1-3 layers of cell wall, we observe the thickness of three layers to be 90 -100 Å, two layers to be 60-70 Å and a single layer to be ~ 30 Å (Figure S1), given that the proposed thickness of PGN in *E. coli* is 20 - 70 Å (Matias et al., 2003; Turner et al., 2013); a single layer was chosen.

**Table 1:** Summary of Simulations.

| OmpA Structure | TolR Structure | BLP | Membrane | Number of atoms | Binds to PGN | Simulation Length (ns) |
| --- | --- | --- | --- | --- | --- | --- |
| None | Open | None | Inner only | 135718 | Yes | 3 X 200 |
| Wildtype | Open | None | Inner and outer | 207787 | Yes | 3 X 200 |
| Wildtype | Open | Yes | Inner and outer |  | Yes | 3 X 200 |
| 1-truncated | Open | Yes | Inner and outer | 238293 | Yes | 3 X 200 |
| Truncated | Open | Yes | Inner and outer | 236933 | Yes | 4 X 200 |
| Truncated | Closed | Yes | Inner and outer | 236789 | No | 3 X 200 |

### Initial binding of open state model of TolR and full length OmpA to peptidoglycan

In simulations of TolR and full length OmpA, both proteins were initially positioned either directly in contact with, or close to the cell wall. Specifically, The OmpA-PGN complex was taken from our previous work and the TolR was positioned with the transmembrane helices embedded in the inner membrane and the periplasmic domain was not in contact with the cell wall(Samsudin, Boags et al. 2017). The shortest distance between the TolR periplasmic domain and the cell wall was around 5 Å at the start of the simulation. This system configuration gives a periplasmic space width of around 170 Å (experimental estimates of the width vary between 100-250 Å (Graham et al., 1991; Vollmer and Seligman, 2010)). The periplasmic domain of TolR in the open conformation was structurally stable in all simulations and showed similar root mean square deviation (RMSD) progressions compared to the C-terminal domain of OmpA (Figure S2). The secondary structure of this domain was also largely preserved during the simulations. In this state TolR has a long unstructured loop in the C-terminus, which remained mobile throughout the simulations.

In all simulations of the wildtype TolR in the open state, binding to PGN was observed. The mechanism of cell wall binding proceeded as follows; the proline (Pro141) residues at the C-terminus of TolR consistently formed the first contact with PGN via its carboxyl group that interacted with either the positively charged amine group of diaminopimelate (m-DAP) or the polar amide and hydroxyl moieties in adjacent sugars (Figure 1 and S3). This was immediately followed by interactions with downstream polar residues (Thr139 and Gln140). The greater flexibility of the unstructured C-terminal loop afforded this initial binding process as these residues were able to ‘snorkel’ towards the PGN. Glu89 and Lys122 found in the more rigid globular domain of TolR strengthened this binding; the former interacted with hydroxyl groups in N-acetyl muramic acid (NAM), whilst the latter formed a salt bridge with carboxyl terminus of m-DAP.

**Figure 1:**
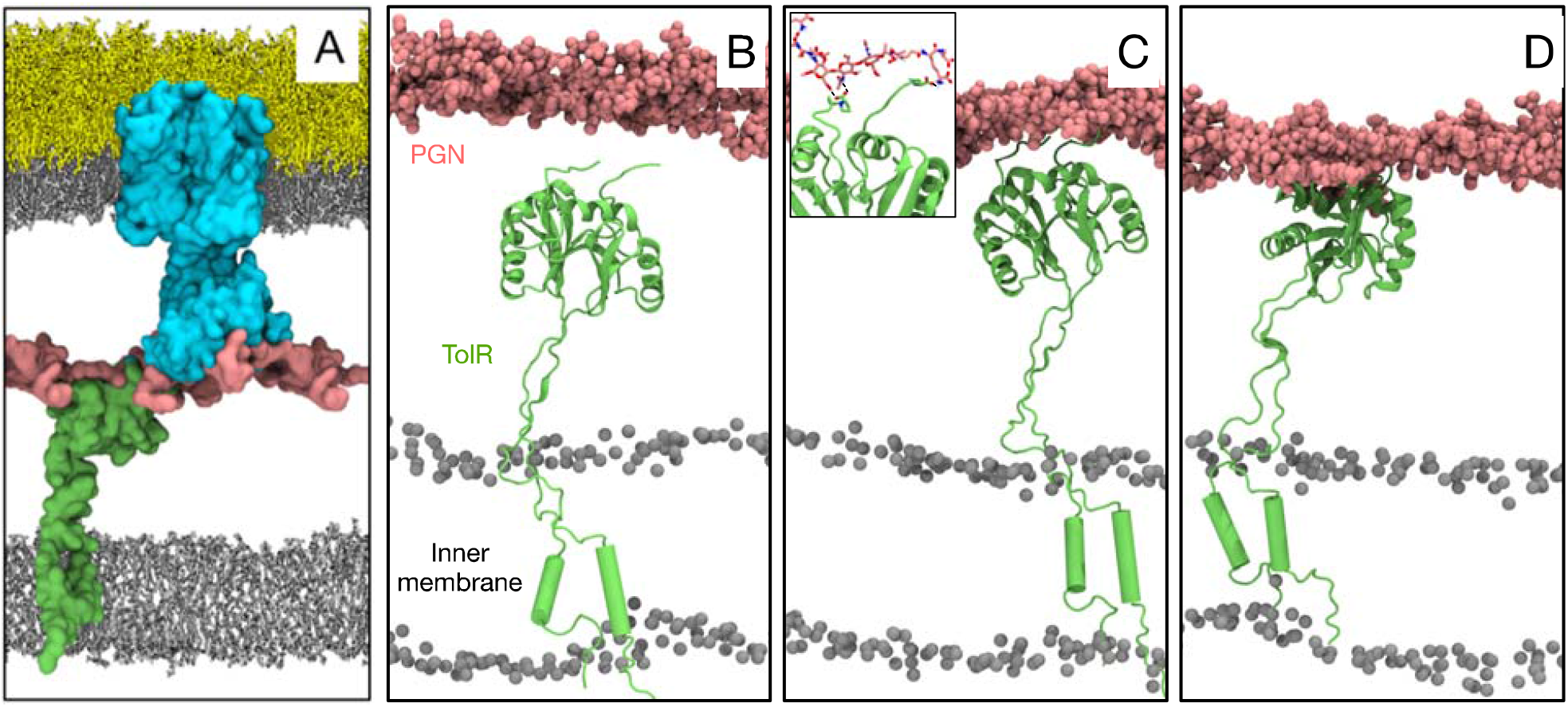
A: Simulation system (OmpA dimer is cyan, PGN is pink, TolR is green, phospholipids are grey and LPS is yellow, water and ions are omitted for clarity). (B-D) Mechanism of TolR binding to the cell wall. (B) TolR in an open state whereby the periplasmic domain is free to bind PGN. (C) The flexible C-terminus of TolR snorkels towards PGN and the carboxyl groups interact with positively charged moieties on PGN. Inset shows example of these interactions (described in detail in Figure 3). (D) The rest of TolR periplasmic domain binds to PGN. The linker between periplasmic domain and N-terminal helices is able to contract to pull the cell wall to the inner membrane.

Within about 10 ns, in each simulation, the TolR linker was extended such that the periplasmic domain was in contact with PGN, in other words PGN binding had occurred. Across all independent repeat simulations, after 200 ns, the cell wall was located about 20 Å (along the z direction, perpendicular to the plane of the membrane) from each protein, reaching a stable position after ~ 100 ns of simulation (Figure 2B). We extended two of these simulations to 500 ns; the binding of OmpA and TolR to the cell wall was maintained (Figure S4). The linker regions of both proteins were only partially extended to enable the cell wall to be maintained at this position. In other words, the proteins had the potential to adopt other arrangements in terms of their location with respect to the cell wall but maintained a position in which the cell was sandwiched equidistant between the two proteins for the duration of these simulations.

**Figure 2:**
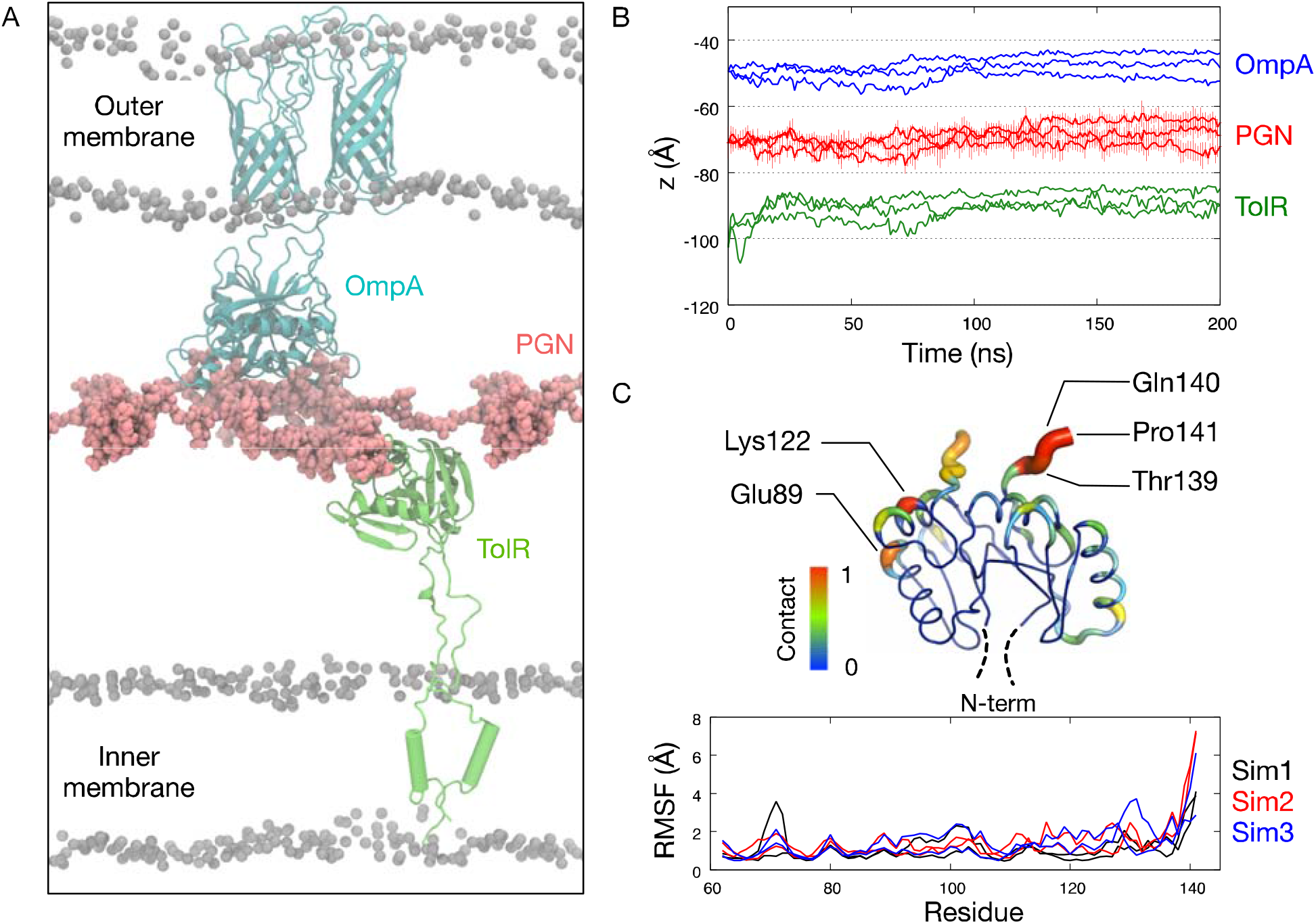
Simulation of TolR with the PGN cell wall and OmpA. (A) A snapshot from the end of a 200 ns simulation. (B) The center of mass motion along the z-axis of OmpA C-terminal domain (residue 189-316), PGN cell wall and TolR periplasmic domain (residue 62-141). Data from three independent simulations are shown. Error bars for the PGN plot indicate standard deviations from three adjacent strands. (C) Root mean square fluctuation (RMSF) of TolR periplasmic domain and the degree of contact each residue made with the cell wall. Two lines in the RMSF plot for each simulation indicate values from two TolR protomers. Contact analysis performed using a distance cut-off of 4 Å and a contact value of 1 indicate one interaction throughout the entire simulations. Residues that made significant contacts are labelled.

Given we have previously reported details of the interactions between OmpA and peptidoglycan (Samsudin, Ortiz-Suarez et al. 2016, Samsudin, Boags et al. 2017), here we focus on the details of the TolR-peptidoglycan interactions. Analysis of the TolR residues in contact with peptidoglycan (where contact is defined as interatomic distance of ≤ 4 Å), reveals that there the five residues that made frequent contacts. These are Glu89, Lys122, Thr139, Gln140 and Pro141 (Figure 2C), they are the same residues identified above as being key to the initial binding and stabilization process. More specifically, the *E. coli* peptidoglycan cell wall has numerous hydroxyl and amide groups on the sugar backbone and the peptide chains that are available for hydrogen bonds (Figure 3A). Additionally, there are three negatively charged carboxyl groups and one positively charged amine group on the non-cross-linked peptide chains (on residue D-glutamate, D-Alanine and diaminopimelate (mDAP)) that can form salt bridges with TolR periplasmic domain. Examples of these interactions involving the key residues identified in Figure 2C are shown in Figure 3B. We found thirteen TolR residues that formed hydrogen bonds with the different parts of the cell wall (Figure 3C). These hydrogen bonds were short-lived with each lasting no longer than 30% of the simulation time scale. Fewer salt bridge interactions were found as there are only five charged residues that are accessible to the cell wall. These salt bridges, however, were longer lasting with lifetime up to 80% of the simulation time scale (Figure 3D). Decomposition of binding free energy shows a larger Coulombic contribution compared to that from VdW interactions, which is concordant with the numerous hydrogen bonds and salt bridges (Figure 3E). Interestingly, the Coulombic contribution of the free energy is correlated to the number of hydrogen bonds formed between TolR and the cell wall, suggesting that hydrogen bonds formation is key for TolR binding (Figure S5). Mapping the electrostatic surface of TolR revealed a predominantly negatively charged surface facing the cell wall contributed by the carboxyl groups on the C-terminus and downstream polar residues (Figure S6). Smaller positively charged patches are found interspersed around this negatively charged surface due to basic residues like Lys122. It makes sense, therefore, that the negatively charged C-terminal region of TolR formed initial binding to the cell wall via electrostatic interactions with positive moieties of the peptidoglycan.

**Figure 3:**
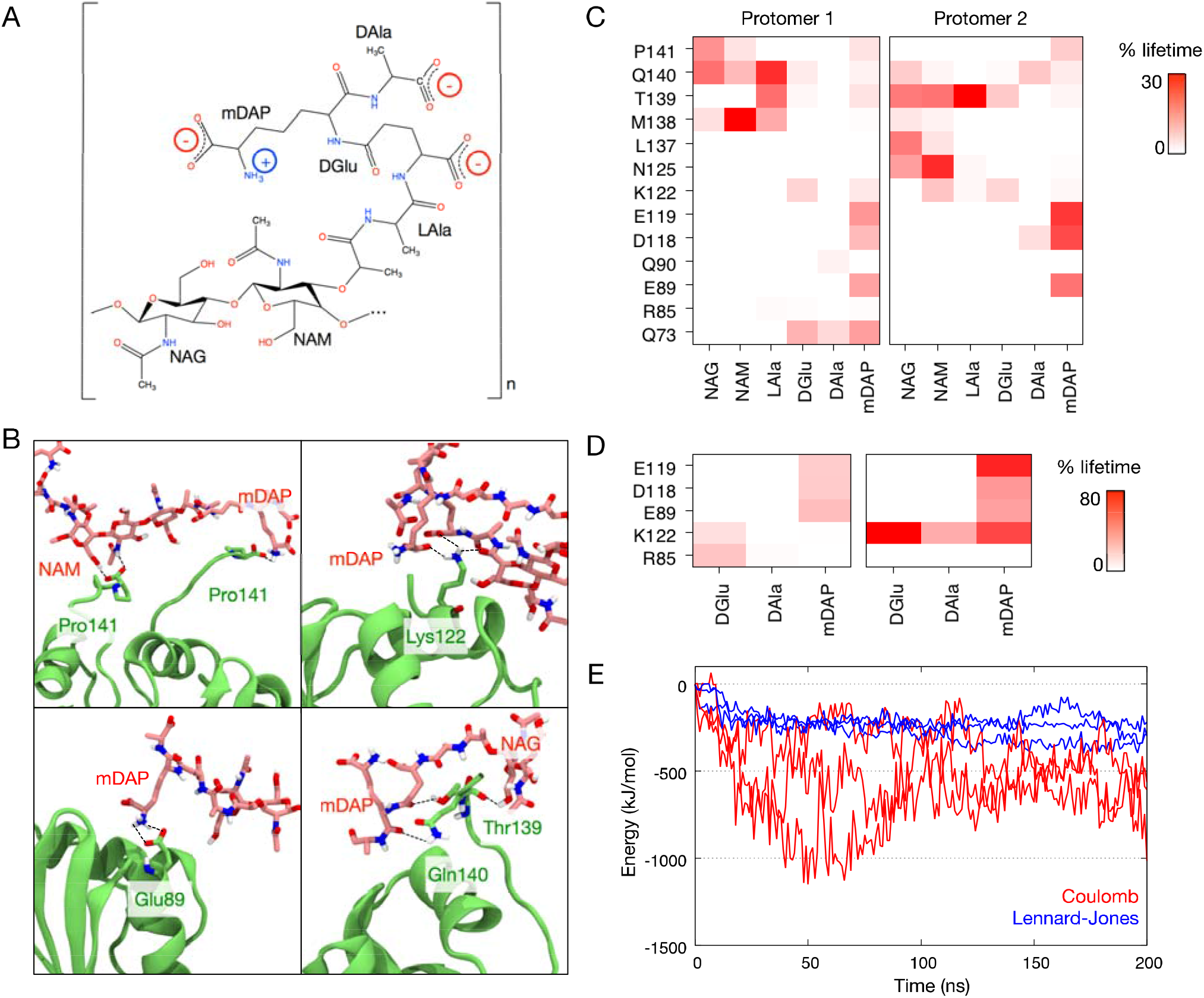
TolR binding to the cell wall is driven by electrostatic interactions. (A) Chemical structure of the peptidoglycan cell wall repeating units with the positions of positive and negative charges highlighted. (B) Four snapshots showing examples of interactions between residues depicted in Figure 1C to diaminopimelate (mDAP), N-acetylmuramic acid (NAM), and N-acetylglucosamine (NAG). (C) Average lifetime of hydrogen bonds formed between residues on both TolR protomers and the different moieties of the cell wall as a percentage of the simulation time. (D) Average lifetime of salt bridges between TolR residues and charged moieties of the cell wall. (E) Binding energy of TolR to the cell wall decomposed into its Coulombic and Lennard-Jones components. Data obtained from three independent simulations.

### TolR-peptiglycan interactions when OmpA is truncated

To determine whether the aforementioned location of the cell wall approximately equidistant from TolR in the inner membrane and OmpA in the outer membrane, is a consequence of the peptidoglycan binding strength of the protein domains being approximately equal, or simply a function of the starting position of the simulations, the C-terminal domain and the linker region of OmpA was truncated, leaving the N-terminal β-barrel (residue 1-172). One BLP trimer was incorporated into these simulations systems to provide an anchor between the cell wall and the outer membrane. When only one of the OmpA protomers was truncated, the cell wall remained bound to both OmpA and TolR in a similar fashion to the wild-type simulations, whilst the BLP tilted to around 60° with respect to the plane of the outer membrane (Figure S7). When both protomers were truncated, however, two distinct behaviours were observed. In three of the simulations TolR remained extended and bound to the cell wall, with the BLP trimer tilted at 60° to enable the location of the cell wall to remain ~ equidistance between the two membranes (Figure 4A). In contrast in one of the four simulations, the linker of TolR contracted such that the bulk of the protein moved to rest on the inner membrane, with BLP almost at right angles to the plane of the outer membrane (Figure 4B). Interestingly, TolR remained bound to peptidoglycan throughout this process. This provides compelling evidence that the peptidoglycan binding domain of TolR proposed by Kleanthous and co-workers does indeed stably bind peptidoglycan, and that this binding can withstand contraction of the TolR linker(Wojdyla, Cutts et al. 2015). Furthermore, these observations show that the balance of non-covalent interactions between proteins in both membranes and the cell wall act like a clamp from both sides in maintaining the position of the cell wall. BLP by itself on the OM side is not sufficient, given its ability to bend and tilt enables significant deviation in the cell wall position. This agrees with experimental studies that showed mutations in either *tolR* or *ompA* genes destabilised the cell envelope, resulting in the formation of outer membrane vesicles in *E. coli* (Deatherage, Lara et al. 2009, Perez-Cruz, Canas et al. 2016).

**Figure 4:**
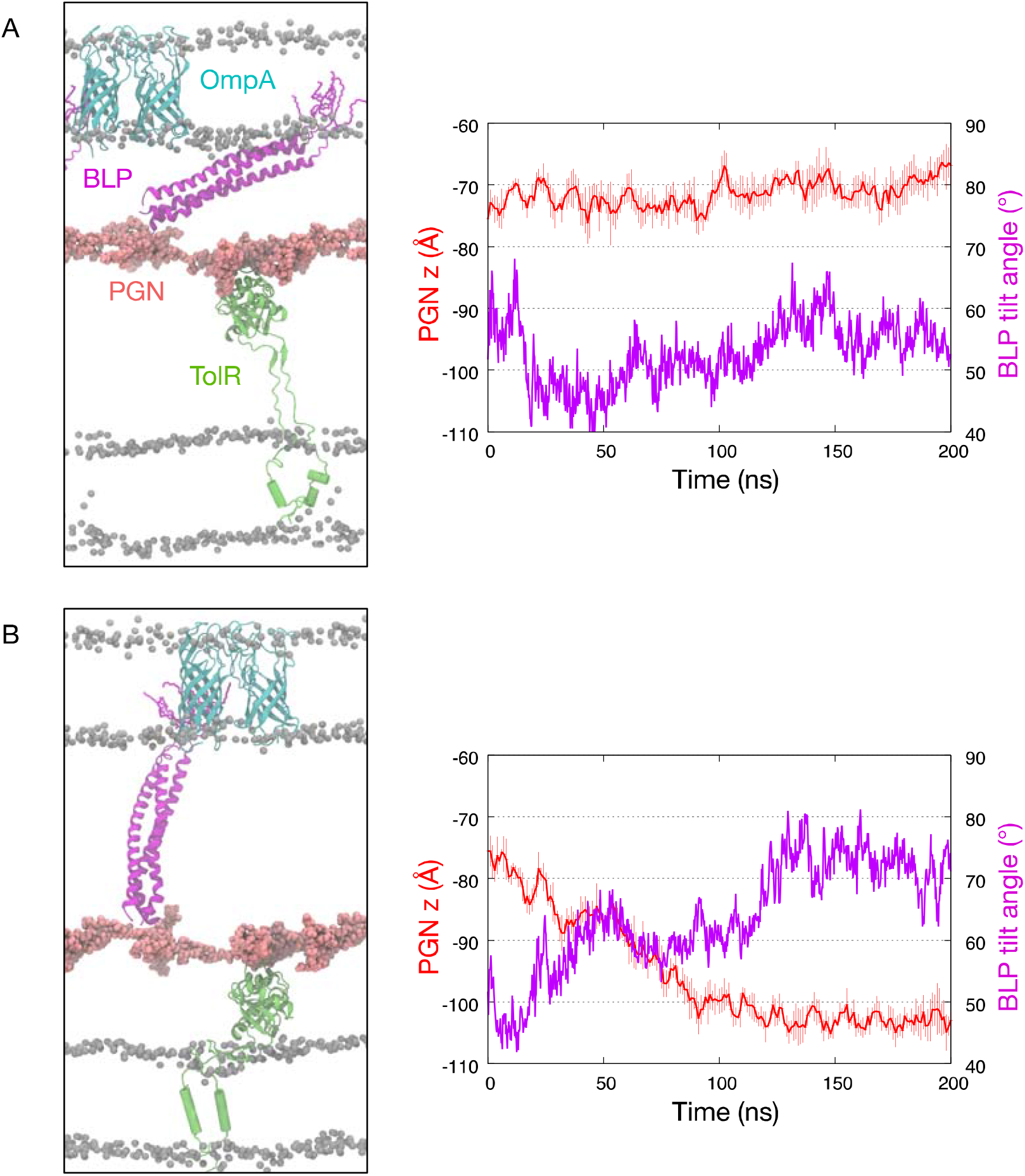
The flexibility of TolR binding to the cell wall. (A) A snapshot from the end of a 200 ns simulation whereby the C-terminal domains and the linker region of OmpA were truncated. In this simulation, the linker between the N-terminal transmembrane domains and periplasmic domains of TolR was fully extended. The graph on the right shows the center of mass motion along the z-axis of the PGN cell wall (red). Error bars indicate standard deviations from three adjacent stands. The angle between the centers of mass of the outer membrane, the N-terminus and the C-terminus of BLP is shown in magenta. (B) A snapshot from a simulation whereby the TolR linker was fully contracted and the TolR bound to the inner membrane, with PGN center of mass motion and BLP tilt angle shown on the right.

### The effect of TolR and OmpA on the structure of the cell wall

Our previous study showed that binding of OmpA to the cell wall caused a local buckling effect on the latter, whereby the surface of the cell wall noticeably curved towards the outer membrane at the point of contact (Samsudin et al., 2017). Similarly here, when only the TolR and the cell wall (no OmpA or BLP) were included in the simulation system, the contraction of the linker pulled the cell wall towards the inner membrane resulting in local curvature of the cell wall. In contrast when both OmpA and TolR were bound to the cell wall, the degree of undulation observed during the simulations was significantly reduced (Figure 5). Interestingly the distortions are also significantly reduced when TolR is bound to the cell wall in the presence of BLP (but without OmpA in the outer membrane). This is in agreement with our previous studies of the outer membrane and the cell wall in which BLP was able to prevent undulations that were otherwise present when OmpA alone was bound to the cell wall. These observations suggest that TolR and OmpA binding to the cell wall from either side of the cell envelope prevent any local distortions caused by either one of them binding alone. From the outer membrane side, BLP also plays a role to this effect, presumably the greater the number of membrane protein interactions with the cell wall, the less distorted the cell wall is, but this hypothesis should be tested with a wider range of peptidoglycan binding proteins.

**Figure 5:**
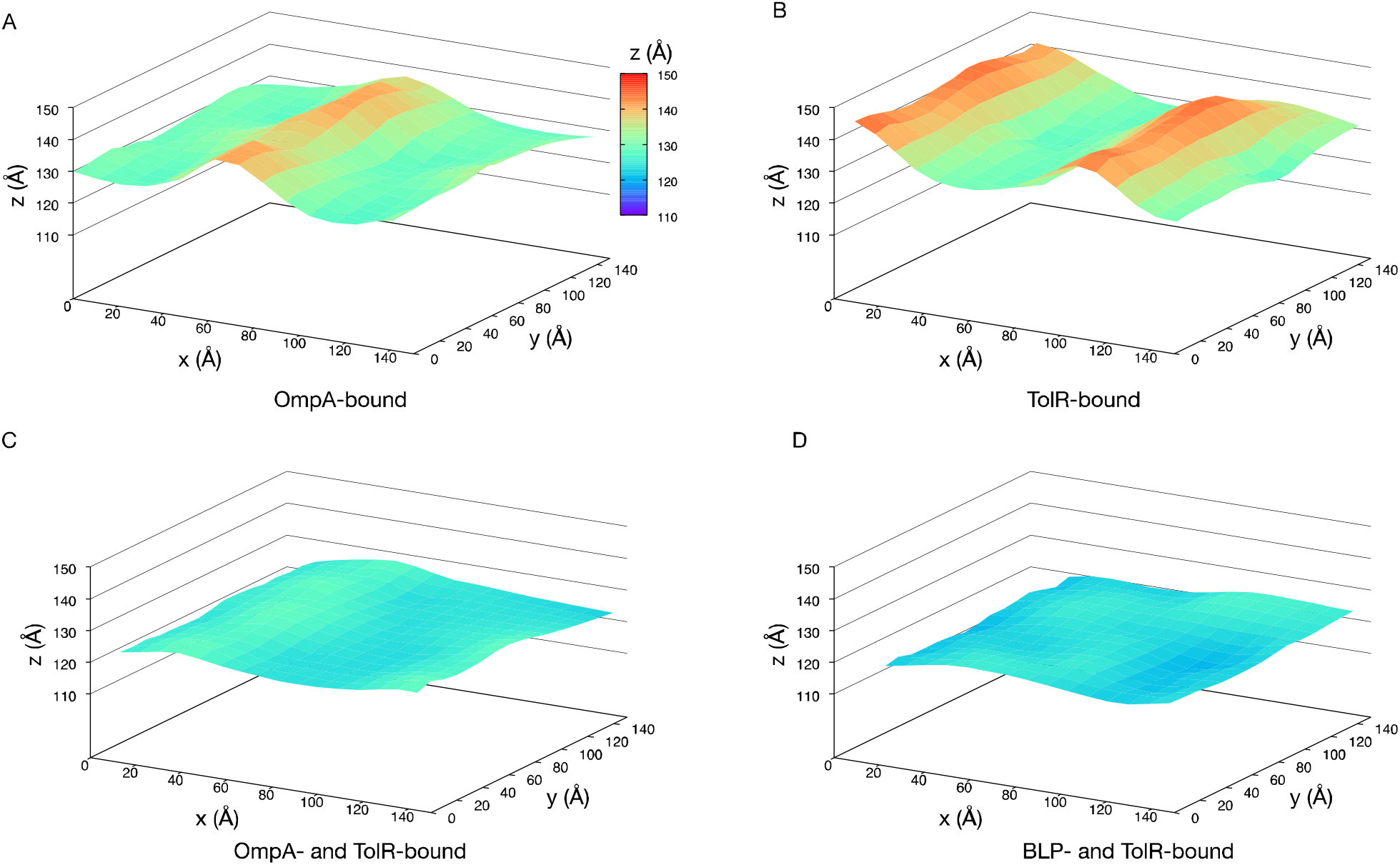
The z-coordinates of PGN cell wall projected into a surface representation. Data taken from (A) the beginning of wild-type simulation whereby only OmpA bound to the cell wall, (B) the end of simulation with only TolR bound to the cell wall, (C) the end of wild-type simulation whereby both OmpA and TolR bound to the cell wall, and (D) the end of truncated OmpA simulation whereby BLP and TolR bound to the cell wall.

### The closed state of TolR does not bind peptidoglycan

As a further test of whether the peptidoglycan binding of TolR is specific to the identified binding domain, or whether other regions of the protein can bind peptidoglycan too, TolR-PGN interactions when TolR is in its closed conformation were also explored. The coordinates for the protein were taken from the X-ray structure (PDB: 5BY4), with the transmembrane helices modelled in as reported by Wojdyla et al. The final snapshot was extracted from our simulation with truncated OmpA dimer in which the TolR linker had contracted to enable interaction of the protein with the inner membrane while still being bound to peptidoglycan. The TolR was replaced by the X-ray structure of the closed state. Thus, at the start of the simulation, BLP was extended and essentially at right angles to the plane of the outer membrane, and the TolR in the closed state was in contact with the inner membrane and within 5 Å from the cell wall (similar to our previous set-up with open state TolR). After 200 ns of simulation BLP tilted to pull the cell wall approximately 20 Å towards the outer membrane and away from TolR (Figure 6). The electrostatic surface potential of the periplasmic domain facing the cell wall in TolR close state is similar to that in the open state, i.e. predominantly negatively charged surface surrounding small positively charged patches (Figure S6). In the closed state, however, the mobile C-terminal domain of TolR responsible for initial interaction with the cell wall, as well as the flexible linker connecting the periplasmic domain and the N-terminal helices are folded together into β-sheet buried within the dimeric structure. The lack of PGN binding of the protein in this state provides further evidence that PGN binding of TolR requires specific domains which are not accessible in the closed state of the protein.

**Figure 6:**
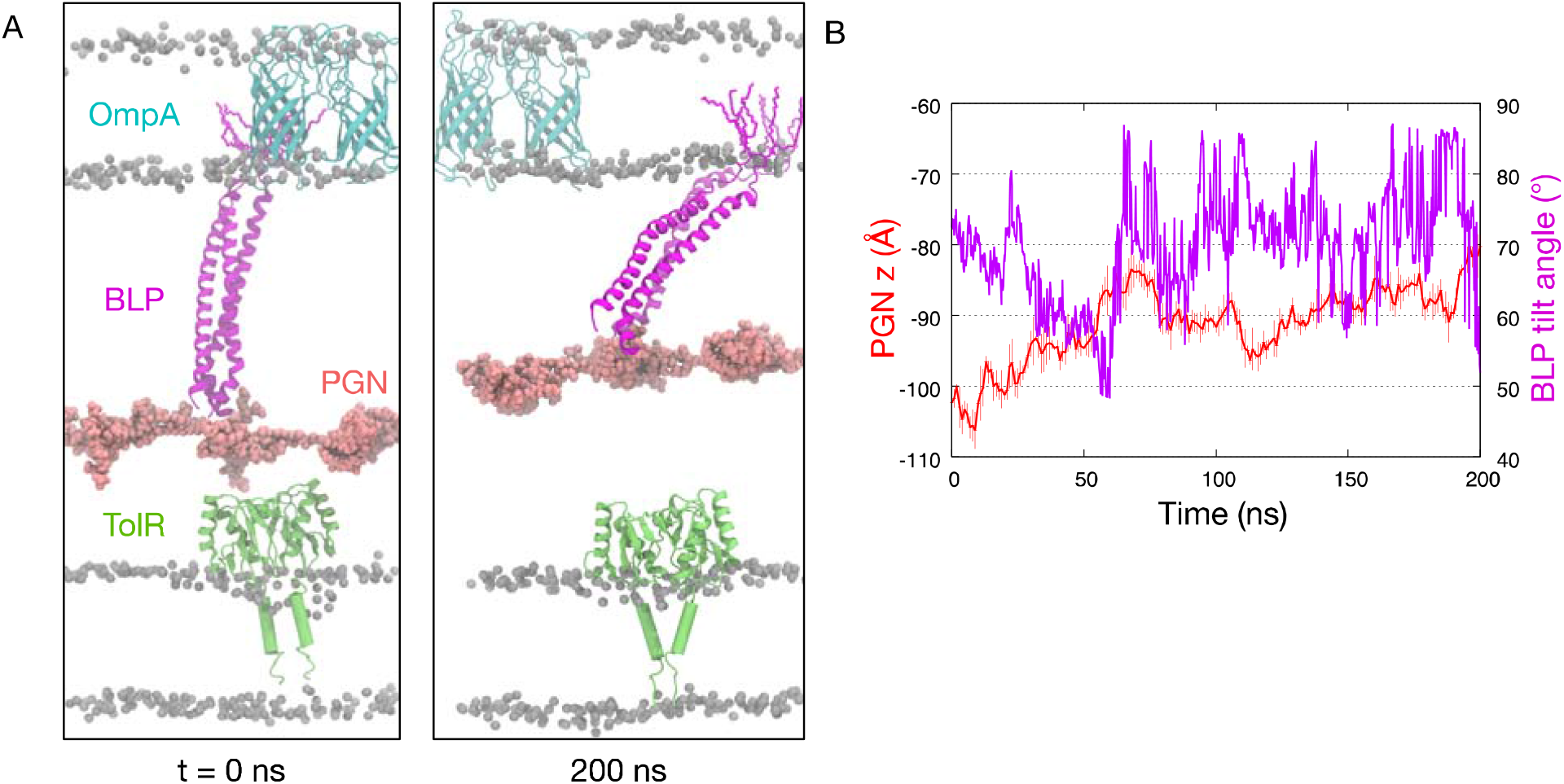
TolR in close conformation does not interact with the cell wall. (A) Snapshot at the beginning (left) and at the end (right) of a 200 ns simulation whereby TolR in its open state was replaced with a close state (PDB: 5BY4). The TolR periplasmic domain did not bind to the PGN cell wall, and the BLP returned to its tilted configuration shifting the cell wall towards the outer membrane. (B) Centre of mass motion along the z-axis for the cell wall and the BLP tilt angle during the simulation as described in Figure 4.

### Conservation of peptidoglycan binding residues of TolR across bacterial species

Having identified Glu89 and Lys122 as key residues in the interaction of TolR and the cell wall, it is worth investigating if this binding mechanism is universally conserved. Sequence alignment of TolR from different Gram-negative bacteria, as well as its structural homolog, ExbD from the TonB system, revealed that both residues are well preserved (Figure S8) (Garcia-Herrero, Peacock et al. 2007). Structural alignment of the open state *E. coli* TolR with *H. influenza* TolR and *E. coli* ExbD lends further support by showing that all of these glutamates and lysines are found in regions that are solvent accessible, and therefore are able to interact with the cell wall.

Our simulations also predicted the role of the highly mobile C-terminal region in providing the first contact with the cell wall via the carboxyl group on Pro141 and two downstream polar residues (Thr139 and Gln140). A similar secondary structure is expected in this region for other homologs. Indeed, the C-terminus of *E. coli* ExbD contains a large unstructured loop, whilst in *H. influenza* TolR the corresponding residues could not be assigned by NMR spectroscopy, which indicate an inherent flexibility. It is worth noting that in the structure of TolR dimer close state, the C-terminus of protomer A via residue 134-138 is a part of a five-stranded β-sheet that includes the N-terminus of protomer B. PMF-dependent unfolding of the N-terminus to form the open state unleashes residue 134-138 to form the flexible region. That these residues are conserved in other bacteria further hints at the presence of an unstructured loop in the open state of TolR in other species. As the first contact made with the cell wall is facilitated by the carboxyl group on the C-terminus instead of its side chain, we conjecture that the specific residue at this position may not be as important as the ability of the entire flexible C-terminal region to ‘snorkel’ towards the peptidoglycan.

## DISCUSSION

In summary, we report the first atomistic molecular dynamics simulation study to our knowledge to include both membranes and the cell wall of *E. coli*. Our simulations enable us to explore the cell wall interactions of the putative open state model of TolR and the impact of these interactions on components inside the periplasm and the outer membrane, namely Braun’s lipoprotein and OmpA respectively. We show that the non-covalent binding of the cell wall of TolR and full length OmpA dimers maintain the location of the cell wall; in other words, the interactions balance each other. TolR binds peptidoglycan via mechanism in which first the terminal residues of each monomer act like arms, snorkeling towards the cell wall and then binding when an appropriate moiety of the peptidoglycan is encountered. Following this, the bulk of the protein is pulled towards the peptidoglycan and then interacts via primarily electrostatic interactions. These interactions are stable over a 500 ns timescale and together with C-terminal of OmpA dimers from the outer membrane side, are able to hold the cell wall at a position that is about equidistant from the two membranes. When the OmpA is truncated to remove its periplasmic, peptidoglycan binding C-terminal domain, TolR is able to ‘pull’ the cell wall towards the inner membrane, even when Braun’s lipoprotein is present. Thus both proteins binding peptidoglycan from both sides are required to hold the cell wall in place. Furthermore when both proteins bind simultaneously, they are able to maintain the cell wall in a flat conformation, when only one protein binds, the cell wall experiences local undulations, here BLP also plays a role in preventing undulations. We provide evidence to show that TolR is not able to bind the cell wall when in its closed state as the PGN-binding is inaccessible. While the emphasis has been on the TolR-peptidoglycan interactions, it is important to note here that here we provide further compelling evidence that Braun’s lipoprotein is able to adjust its tilt angle to accommodate a range of outer membrane to cell wall distances. We show that for fine control over cell wall position, Braun’s lipoprotein alone is not sufficient; the non-covalent interactions from proteins in both membranes play a crucial role. This is in line with experimental observations that *E. coli* mutants missing with OmpA or TolR produce more vesicles than normal cells, presumably at least in part due to the easier detachment from the cell wall. Overall a picture is beginning to build up that suggests the local conformation of the cell wall is dependent upon the non-covalent interactions from proteins in both membranes. It is important to note here that other proteins that bind to the cell wall from both sides, such as PAL, and MotB are not considered in the present study, these too will undoubtedly impact on properties such undulations/distortion and overall position of the cell wall with respect to the two membranes (Parsons, Lin et al. 2006, Roujeinikova 2008). It is possible and likely, based on our current findings, that the cell wall location varies as a function of the local protein composition of both membranes, and therefore one would expect an undulating structure at the molecular level of detail. Further studies with additional inner and outer membrane proteins and varying copy numbers of BLP are needed to explore these aspects. Our current study provides an important step towards the study of the mechanical interplay between the two membranes and cell wall of Gram-negative bacteria at the atomistic level.

## EXPERIMENTAL PROCEDURES

### Protein and Cell Wall Parameters

The full-length OmpA dimer model was obtained from Carol Robinson (Marcoux et al., 2014); their structural stability in a model OM has been verified in our previous work (Ortiz-Suarez et al., 2016). The model of the TolR dimer in the open state was obtained from Phillip Stansfeld (Wojdyla, Cutts et al. 2015). A PGN network consisting of three strands of 10 repeating NAG-NAM-peptide units was constructed and positioned ~90 Å from the surface of the lower leaflet of the OM. The BLP homotrimer was built based on the structure from Shu et al. (PDB: 1EQ7) (Shu et al., 2000) with the last residues on both the N- and C termini manually added back using PyMOL (DeLano, 2002). The N-terminus was in turn attached to the tripalmitoyl-S-glyceryl-cysteine residues to incorporate the BLP into the inner leaflet of the OM. The parameters for tripalmitoyl-S-glyceryl-cysteine were constructed from the standard GROMOS 54A7 force field with the GROMOS 53A6OXY ether parameters used for the linkage region (Horta, Fuchs et al. 2011). PGN was then covalently linked to the lysine on one of the C termini of the BLP trimer via its m-DAP residue. The linkage was constructed using the standard GROMOS 54A7 parameters.

### Outer and Inner Membrane Construction

The OM model was asymmetric: the upper leaflet was made entirely of full-rough Ra LPS lipids of the R1 core type, whereas the lower leaflet comprised a mixture of phospholipids in the following ratio; 90% 1-palmitoyl 2-cis-vaccenic phosphatidylethanolamine, 5% 1-palmitoyl 2-cis-vaccenic phosphatidylglycerol, and 5% 1-palmitoyl 2-cis-vaccenic 3-palmitoyl 4-cis-vaccenic diphosphatidylglycerol, known as cardiolipin. This OM model has been validated in our previously reported studies (Samsudin, Boags et al. 2017).

The IM model was symmetric: both leaflets were made entirely of phospholipid in the following ratio; 90% 1-palmitoyl 2-cis-vaccenic phosphatidylethanolamine, 5% 1-palmitoyl 2-cis-vaccenic phosphatidylglycerol, and 5% 1-palmitoyl 2-cis-vaccenic 3-palmitoyl 4-cis-vaccenic diphosphatidylglycerol, this bilayer has the same composition per leaflet as the inner leaflet of the OM model.

### Simulation Systems

Simulations systems were constructed with full length dimeric OmpA in the OM and the extended conformation of TolR in the IM. OmpA was inserted into the OM using the membed protocol from the GROMACS package (Wolf et al., 2010). TolR was inserted into the IM manually and any overlapping lipids were removed. Both wild type proteins were simulated with a starting point (distance) to the cell wall for OmpA and (distance) for TolR, where the proteins can either interact or move away from the PGN structure. Truncation of OmpA C-terminal domain was performed, whereby residue 173-316 was removed from either one of the protomers or from both. In the truncated systems BLP is included as previously described. The cell wall was included in all simulation systems, where the cross linking was parameterized from the peptide bond parameters from the GROMOS 54A7 force field (Schmid, Eichenberger et al. 2011). This cross linking is periodic, where the sheet is self-bonded at the end of the strands, using the existing glycan parameters from GROMOS 54A7, allowing the sheet to behave as an infinite layer, we have described and used this approach in our previous studies (Samsudin, Boags et al. 2017).

All simulation systems are run at a neutral charge with a concentration of 0.2 M sodium chloride ions in solution. All repeats of the systems are run from the same starting point, where the independent repeats are achieved by regenerating the velocities of the atoms randomly at the start point for each repeat.

### Simulation Protocols

All simulations were performed using the GROMACS 2018 simulation package, the GROMOS 54A7 force field with the SPC water model (Berendsen, Postma et al. 1981, Schmid, Eichenberger et al. 2011, Abraham, Murtola et al. 2015). Each simulation was run for 200 ns, and at least three independent repeats of each simulation was performed, giving at least 600 ns for each system simulated. Temperatures of 310 K were maintained using the velocity rescale thermostat using a time constant of 1 ps (Bussi et al., 2007). The pressure was maintained semi-isotropically at 1 atm using the Parrinello-Rahman barostat with a time constant of 1 ps (Parrinello and Rahman 1981). All bonds were constrained using the LINCS algorithm to allow for an integration time step of 2 fs (Hess, Bekker et al. 1997). Long-range electrostatics were described using the particle mesh Ewald method (Essmann et al., 1995). The short-range electrostatic cutoff used was 1.4 nm and the short-range van der Waals cutoff was also 1.4 nm.

### Sequence and structural alignment and electrostatic surface analysis

All sequences were fetched from the Uniprot webserver and aligned in JalView (Waterhouse, Procter et al. 2009) using the ProbCons algorithm(Do, Mahabhashyam et al. 2005). The structure of TolR open state model was aligned to E. coli ExbD (PDB: 2PFU)(Garcia-Herrero, Peacock et al. 2007) and H. influenza TolR (2JWK) (Parsons, Grishaev et al. 2008) using PyMOL(Schrodinger). Electrostatic profile of TolR was calculated using APBS (Baker, Sept et al. 2001).

## SUPPLEMENTAL INFORMATION

Supplemental Information includes Figures S1-8 and can be found with this article at:XXX

## ACKNOWLEDGEMENTS

This research was supported by UK Biotechnology and Biological Sciences Research Council (Grant No. BB/M029573/1) and the Institute for Life Sciences, University of Southampton. This project made use of time on Iridis V supercomputers provided by the University of Southampton and on ARCHER provided by HECBioSim through EPRSC (Grant No. EP/L000253/1). The authors thank Phillip Stansfeld for providing the coordinates of the TolR open state model and Colin Kleanthous for useful discussions.

